# P-bodies and the miRNA pathway regulate translational repression of *bicoid* mRNA during *Drosophila melanogaster* oogenesis

**DOI:** 10.1101/283630

**Authors:** John M. McLaughlin, Daniel F.Q. Smith, Irina E. Catrina, Diana P. Bratu

## Abstract

Embryonic axis patterning in *Drosophila melanogaster* is partly achieved by mRNAs that are maternally localized to the oocyte; the spatio-temporal regulation of these transcripts’ stability and translation is a characteristic feature of oogenesis. While protein regulatory factors are necessary for the translational regulation of some maternal transcripts (e.g. *oskar* and *gurken*), small RNA pathways are also known to regulate mRNA stability and translation in eukaryotes. MicroRNAs (miRNAs) are small RNA regulators of gene expression, widely conserved throughout eukaryotic genomes and essential for animal development. The main *D. melanogaster* anterior determinant, *bicoid*, is maternally transcribed, but it is not translated until early embryogenesis. We investigated the possibility that its translational repression during oogenesis is mediated by miRNA activity. We found that the *bicoid* 3’UTR contains a highly conserved, predicted binding site for miR-305. Our studies reveal that miR-305 regulates the translation of a reporter gene containing the *bicoid* 3’UTR in cell culture, and that miR-305 only partially contributes to *bicoid* mRNA translational repression during oogenesis. We also found that Processing bodies (P-bodies) in the egg chamber may play a role in stabilizing *bicoid* and other maternal transcripts. Here, we offer insights into the possible role of P-bodies and the miRNA pathway in the translational repression of *bicoid* mRNA during oogenesis.

## INTRODUCTION

The genes and signaling pathways that control embryonic axis patterning in *Drosophila melanogaster* have been studied for several decades. Valuable insight gained from this model system includes the importance of mRNA localization and the regulation of mRNA stability and translation for development. The *D. melanogaster* egg chamber develops as a multicellular structure containing the germline, one oocyte and 15 support cells (nurse cells), as well as somatic cells (follicle cells) (1). The oocyte nucleus is arrested in meiosis for the majority of egg chamber development, therefore gene products are provided by the adjacent nurse cells via intercellular ring canals; the microtubule (MT) network mediates most of this long-range transport.

The main patterning determinants of *D. melanogaster, gurken* (*grk*)*, nanos* (*nos*)*, oskar* (*osk*) and *bicoid* (*bcd*), are localized as mRNAs to discrete compartments of the oocyte by mid-oogenesis (1). During oogenesis, *gurken* and *nanos* mRNAs localize to the dorsoventral and posterior compartments of the oocyte, respectively. When translated, their encoded proteins act to specify the dorsoventral and anteroposterior axes of the future embryo. The *bcd* mRNA is localized to the anterior of the oocyte during oogenesis and translated following egg activation, at which point the resulting Bicoid transcription factor activates anterior patterns of gene expression in the developing embryo (2). Protein factors that bind mRNA *cis*-elements are responsible for mediating the localization and translational control of several of these patterning transcripts (3). For instance, the *nos* 3’UTR contains a 90-nucleotide region termed the translational control element (TCE). Stem loops within the TCE are bound by two protein factors, Glorund and Smaug, which act to translationally repress unlocalized *nos* mRNA in the oocyte and embryo (3).

An additional layer of mRNA regulatory control is provided by Processing bodies (P-bodies): cytoplasmic, non-membrane bound organelles that play a role in the storage and/or decay of cellular mRNA. In *D. melanogaster*, P-bodies contain proteins involved in different aspects of RNA metabolism, including the mRNA decapping enzymes Dcp1/2, decapping activators Me31B and Trailer Hitch (Tral), and the 5’ to 3’ exonuclease Pacman (4). Previous studies have implicated P-bodies in the post-transcriptional regulation of *D. melanogaster* maternal mRNAs; ovarian P-bodies harbor *grk*, *osk*, and *bcd* mRNAs (5, 6), and the major P-body component Me31B is also known to play a role in the repression of *osk* translation (5). Interestingly, in mammalian cells, P-bodies are involved in the functioning of microRNAs (miRNA), a small RNA class of post-transcriptional regulators of mRNA stability and translational repression (7, 8).

miRNAs have gained attention in the past decade as a widespread class of small regulatory RNAs; they are now known to be essential regulators of developmental timing, differentiation, and morphogenesis in most animals. A mature miRNA, in complex with an Argonaute protein (Ago1 in *Drosophila*), is guided via base pairing to the 3’UTR of an mRNA target, to effect translational silencing or RNA decay. Less is known regarding miRNAs and their targets during *D. melanogaster* oogenesis compared to other tissues. Global miRNA function is required for germline stem cell maintenance, indicating early essential functions in egg chamber development (9–11). Null alleles of core miRNA biogenesis factors, such as *dicer-1* and *ago1*, are associated with early developmental arrest in the female germline. MiRNA gene mutants have been lacking in *D. melanogaster*, however a recent study from the Stephen Cohen group (12), which generated several dozen targeted miRNA knock-outs, will enable further studies of the functions of individual miRNAs during fly development. To date, several publications have demonstrated roles for specific miRNAs in various aspects of egg chamber patterning and early embryonic development (13–15). For example, miR-184 acts during egg chamber development to regulate stem cell differentiation, dorsal appendage patterning, and pair rule gene activation during embryogenesis (16), and miR-318 regulates follicle cell gene amplification and chorion formation (17).

Previous work on *bcd* mRNA localization suggests that its translational control is mediated by the 3’UTR, which is a potential site of regulation by miRNAs. The *bcd* 3’UTR contains a Nanos Response Element (NRE), an RNA sequence motif bound by the translational repressor Pumilio (Pum). The NRE has been previously shown to regulate *bcd* mRNA’s stability during early embryogenesis (18), but the possible role it plays during oogenesis has not yet been examined. Heterologous reporter constructs bearing the *bcd* 3’UTR display the endogenous *bcd* mRNA’s translational timing. However, an *osk* mRNA bearing the *bcd* 3’UTR, in which the NRE is deleted, is translated at the oocyte’s anterior during oogenesis (19). Additional observations suggest that the *bcd* NRE is involved in its translation control. Ectopic expression of Nanos at the oocyte anterior prevents the translation of *bcd* mRNA during embryogenesis, and this effect is dependent on the NRE (20). Furthermore, the *bcd* 5’UTR is dispensable for its translational repression during oogenesis (21), underscoring the idea that its translational timing is likely mediated by its 3’UTR.

Because *D. melanogaster* miRNAs mainly act on a target’s 3’UTR, we examined the possibility that one or more miRNAs contribute to the translational repression of *bcd* mRNA during oogenesis. Using computational methods, we identified miR-305 as a possible regulator of *bcd* mRNA translation, we confirmed miR-305 expression in ovaries, as well as demonstrated its ability to repress translation of a luciferase reporter gene bearing the *bcd* 3’UTR in a cell culture assay. Moreover, via knock-down of individual P-body components in the egg chamber, we examined the relationship between the function of ovarian P-bodies and the expression levels of *bcd* and other maternal transcripts during oogenesis.

## MATERIALS AND METHODS

### Cloning

#### UASp*-*gfp-bcd

A plasmid containing the complete *bcd* locus bearing an N-terminal GFP tag was a gift from Thomas Gregor (22). Primers spanning the *bcd* 5’UTR and ∼1 kb downstream of the annotated 3’UTR were used to amplify the *bcd* fragment that was cloned into the pENTR/D-TOPO vector (Gateway System, Invitrogen). This transgene was then recombined into a modified UASp vector (gift from Jennifer Zallen) to produce *UASp-gfp-bcd* for transformation into *D. melanogaster*.

#### gfp-bcd

Primers spanning 200 nucleotides (nt) of the *bcd* promoter sequence and 1 kilobase pairs (kb) downstream of the annotated 3’ UTR were used to amplify the *bcd* fragment that was cloned into pBID-G (Addgene, catalogue #35195), a *D. melanogaster* transgene expression vector.

#### pSiCheck-*bcd* 3’UTR

The full *bcd* 3’UTR was PCR amplified from a Bcd-GFP plasmid (gift from Thomas Gregor lab) (22) and cloned into Xho1 and Not1 sites of the pSiCheck 2 vector (Promega). Mutagenesis to generate each of the *bcd*-3’UTR reporters was performed by PCR using mismatched primers (S4 table; Primer Sequences) and verified by sequencing.

### Transgenic fly stocks

‘*UASp*-*gfp-bcd*’, ‘*UASp*-*gfp-bcd* seed mutant’, and *‘gfp-bcd’* fly stocks were generated using the Phi C31 integrase system. The *UASp-gfp-bcd* and *UASp-gfp-bcd* seed mutant transgenes were inserted into an attP40 landing site on chromosome 2; the ‘*gfp-bcd’* transgene was inserted into an attP18 landing site on the X chromosome (Genetic Services Inc). miR-305 KO and miR-305 sensor fly lines were gifts from Stephen Cohen (23).

### Commercial fly stocks

The following fly stocks were obtained from the Bloomington *Drosophila* Stock Center:

*cuc^1^* (BL #11765), Df(2L)BSC190 (BL #9617),

maternal Gal4 driver w*; P[matα-Gal4-VP16]V37 (BL #7063), and Gal4-inducible RNAi TRiP lines

*mCherry* (BL #35785), *ago1* (BL #33727), *tral* (BL #38908), *me31b* (BL #38923), *gw182* (BL #34796),

*pacman* (BL #34690), *ccr4/twin* (BL #32490), *not1* (BL #32836), *pop2* (BL #52947),

*thor/4E-BPT* (BL #36815) and *staufen* (BL #43187). Fluorescently-tagged fly lines were obtained from the Kyoto *Drosophila* Stock Center: *me31b-yfp* (#115460), *tral-gfp* (#115090) and *pum-gfp* (#115589).

### miRNA primer design

Primers for amplification of specific mature miRNAs were designed as previously described (24). Primer sequences for miRNA amplification, cloning, and mutagenesis are included in S4 Table (Primer Sequences).

### RNA isolation

For small RNA isolation, ovaries were dissected into cold 1X PBS and kept on ice, then washed with cold 1X PBS, and RNA isolated using the PureLink miRNA Isolation Kit (ThermoFisher) according to the manufacturer’s instructions. For total RNA isolation, ovaries were dissected into cold 1X PBS and kept on ice, then washed with cold 1X PBS, and RNA isolated using TRIzol (ThermoFisher) according to the manufacturer’s instructions.

### RT-qPCR

Reverse transcription reactions were performed as follows: for miRNAs, 100 ng of small RNA was incubated with 2 pM RT primer at 70 °C for 5 minutes, cooled on ice for 5 minutes, followed by addition of GoScript RT enzyme (Promega) and incubation at 42 °C for 1 hour. For cDNA synthesis of mRNA targets, the protocol was identical except that 250 ng of total RNA was incubated with 0.5 μg of dT18-20 or random hexamers. qPCR reactions were performed in a Roche Lightcycler 480 (Roche Molecular Systems, Inc.). Each PCR reaction contained 1 μl of cDNA from RT reaction, 2 μl primer solution (10 μM of each, forward and reverse primers), 2 μl dH2O, and 5 μl SYBR Green enzyme mix (Roche Molecular Systems, Inc); 95 °C denaturation for 5 minutes, followed by 40 cycles of 95 °C for 20 seconds, 58 °C for 15 seconds, 58 °C for 15 seconds.

### Luciferase assays

Luciferase assays were performed using the Promega Dual-Glo assay as previously described (25). The pSiCheck 2 vector contains a *Renilla* luciferase gene placed under the control of a 3’UTR of interest and an independent firefly luciferase gene that serves as an internal control for transfection efficiency. Briefly, the DsRed-miRNA and pSiCheck constructs were co-transfected into *D. melanogaster* S2 cells using Effectene transfection reagent (QIAGEN) and plated at a density of 1.1-1.2 × 10^6^ cells/ml. Cells were incubated for three days at room temperature, lysed, and both *Renilla* and firefly luciferase levels were measured using a Veritas plate luminometer (Turner Biosystems).

### Data analysis

For luciferase reporter assays, GraphPad Prism 6 was used to perform two-tailed student’s *t*-tests and generate box plots. For RT-qPCR experiments, Ct values were determined using the Roche LightCycler 480 software and exported to Microsoft Excel to calculate mean values and perform two-tailed student’s *t*-tests. For quantitation of GFP expression in *UASp-gfp-bcd* egg chambers, pixel values were determined using ImageJ (Fiji platform) (26) and then exported to Microsoft Excel to calculate mean values.

### Single-molecule fluorescence *in situ* hybridization (smFISH)

Ovaries were dissected into 1X PBS, fixed with 4% paraformaldehyde in 1X PBS, then washed with 1X PBS before performing smFISH as previously described (27). Briefy, following fixation ovaries were pre-hybridized in wash buffer, incubated with Cy5 labeled *bcd* mRNA specific smFISH probe solution overnight at 37 °C, then washed in wash buffer before being mounted on a glass slide with Prolong Gold Media (ThermoFisher).

### Immunofluorescence

Dissected ovaries were fixed with 4% paraformaldehyde in 1X PBS, permeabilized in 1% Triton X-100, 1% BSA in 1X PBS for 2 hours, and washed with 1X PBS. Egg chambers were incubated overnight at room temperature or 4 °C with rabbit anti-Bicoid (1:30) in 0.3% Triton X-100, 0.1% BSA in 1X PBS, washed with 0.05% Triton X-100, 0.1% BSA in 1X PBS, followed by incubation with a fluorescently labeled secondary antibody at 1:1000 dilution for at least 2 hours at room temperature. Ovaries were washed with 0.05% Triton X-100 in 1X PBS and mounted with Prolong Gold Media (ThermoFisher). Anti-Bicoid is a rabbit polyclonal antibody obtained from Santa Cruz Biotechnology (product number d-200), and raised against amino acids 295-494 in *Drosophila* Bicoid.

### Imaging

Imaging was performed with a LeicaDMI-4000B inverted microscope (Leica Microsystems) mounted on a TMC isolation platform (Technical Manufacturing Corporation), with a Yokogawa CSU 10 spinning disc head and Hamamatsu C9100-13 ImagEM EMCCD camera. This set up includes 491, 561, and 638 nm diode lasers. Images were acquired with 40X/NA = 1.25 or 63X/NA = 1.4 oil objectives, using Volocity software (PerkinElmer). Z-stacks were acquired using a manual XY-stage with piezo-Z. Image processing and analysis were performed using ImageJ software.

## RESULTS

### *bicoid* mRNA localizes to P-bodies during oogenesis

A previous immuno-EM study of maternal mRNA localization in P-bodies demonstrated that *bcd* mRNA is enriched in the P-body “core” relative to another maternal transcript, *grk*, which is detected primarily at the P-body periphery (6). However, these analyses were limited to P-bodies located at the anterior cortex of the oocyte. To determine the extent of *bcd* mRNA and P-body marker(s) co-localization throughout the whole egg chamber, we used transgenic lines expressing protein-trapped Me31B-YFP or Tral-GFP. We visualized *bcd* mRNA by smFISH in fixed ovaries, and observed that most *bcd* mRNA particles are present in Me31B and Tral foci, in both the nurse cell and oocyte compartments of the egg chamber during early (Fig 1A) mid and late oogenesis (Fig 1B).

**Fig 1.**
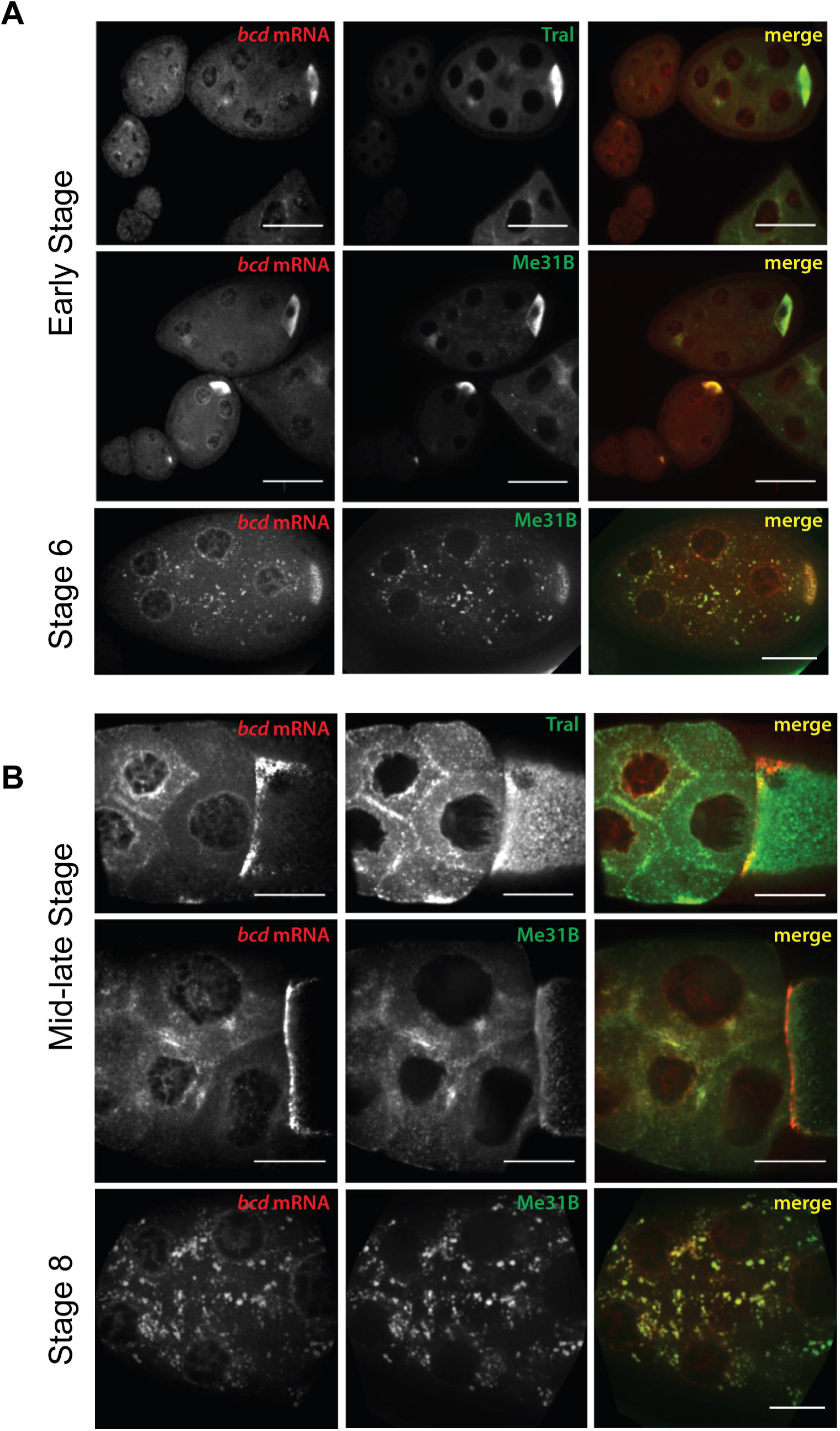
Accumulation of *bicoid* mRNA in egg chamber P-bodies. (A) Early and (B) mid-to-late stage Me31B-YFP and Tral-GFP (green) egg chambers were probed using smFISH for *bcd* mRNA (Cy5-labeled Stellaris probes: red). The merge panels show areas where *bcd* mRNA co-localizes with these P-body components (yellow). Images were acquired with 40X/NA = 1.25 oil objective, and are maximum intensity XY projections of 4 optical Z slices (Z step of 0.5 μm). Scale bar = 25 μm. Representative images were selected from at least 3 independent experiments.

### The mRNA deadenylase pathway is required for P-body formation in egg chambers

P-bodies store mRNAs for translational repression or decay, and both processes require a shortened poly-A tail. A functional mRNA deadenylase pathway is essential for P-body formation in yeast and mammalian cells (28). To investigate if the same requirements apply during *D. melanogaster* oogenesis, we visualized P-body markers Me31B-YFP and Tral-GFP in egg chambers depleted of members of the mRNA deadenylase and 5’->3’ exonuclease pathways: the 5’ to 3’ exonuclease Pacman, the exonuclease subunits Ccr4 and Pop2, and the scaffolding subunit Not1.

In *pacman* knock-down egg chambers, P-bodies seemed to increase in size, consistent with previous findings from Pacman-null egg chambers (Fig 2B vs. 2A) (4). In contrast, depletion of Not1 resulted in a substantial loss of detectable P-bodies (Tral), with Me31B signal accumulated at the nurse cell periphery, suggesting a critical role for this scaffolding subunit in P-body assembly and cytoplasmic localization (Fig 2D vs. 2A). A similar loss of P-bodies and *bcd* mRNA was observed in the *pop2* (Fig 2E vs 2A; Tral), but not in the *ccr4* knock-down (Fig 2C vs. 2A), suggesting that Pop2 may be the more active exonucleolytic subunit contributing to P-body function in egg chambers.

**Fig 2.**
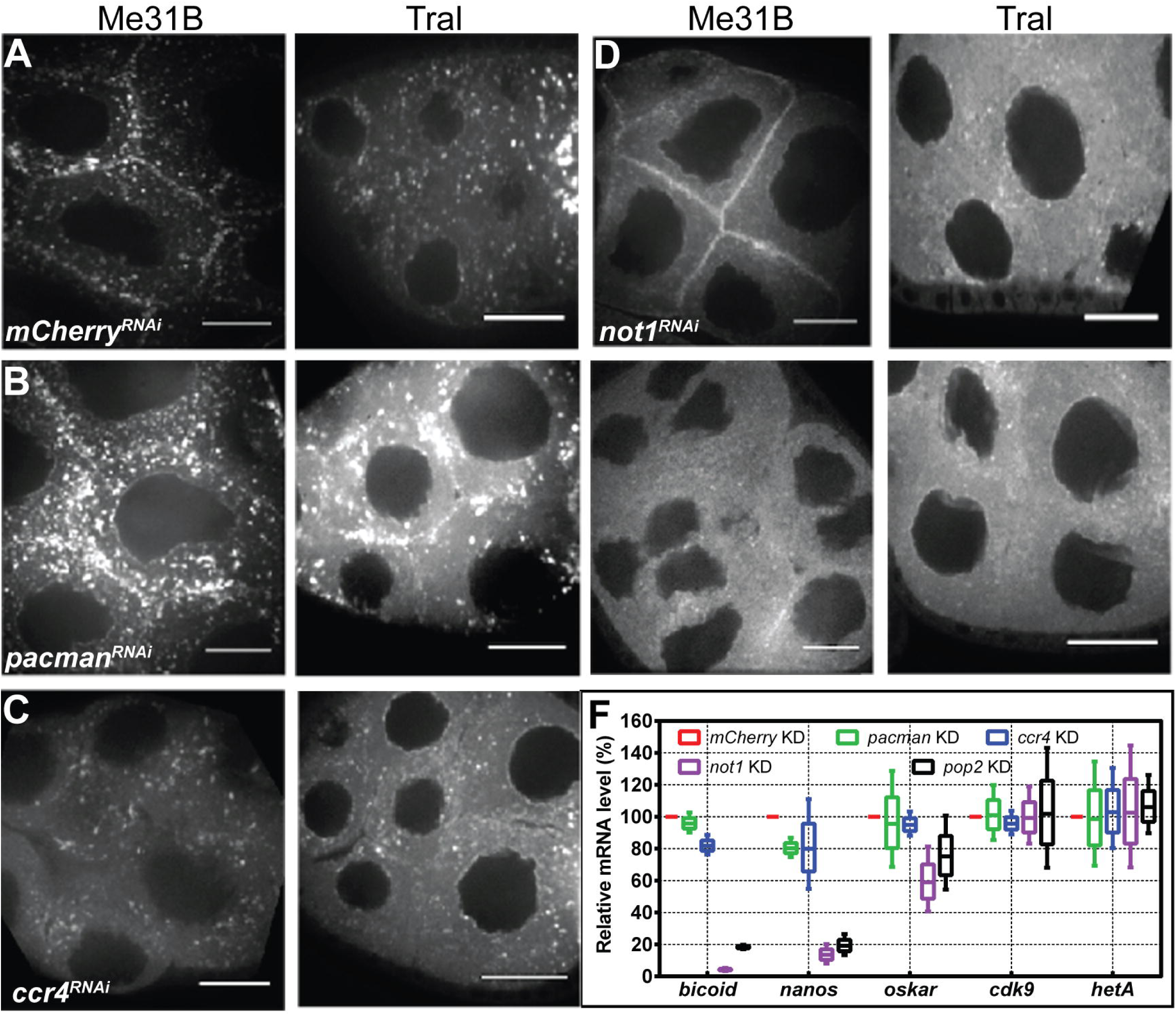
Reduced levels of P-body components are directly correlated with reduced maternal mRNA levels. Gal4-inducible RNAi transgenes targeting members of the CCR4-Not deadenylase and RNA decay pathways were driven in Me31B-YFP and Tral-GFP egg chambers. Ovaries isolated from the indicated knock-downs were fixed and imaged to analyze the distribution of P-body marker proteins. (A) Control (mCherry) and (B-E) P-body component knock-downs in Me31B-YFP or Tral-GFP egg chambers, as indicated. Images were acquired with a 63X/NA = 1.4 oil objective and are maximum intensity XY projections of 4 optical Z slices (Z step of 0.5 μm). Scale bar = 25 μm. Representative images were selected from at least three independent experiments. (F) Total RNA was isolated from whole ovaries dissected from the control (red) and *pacman* (green), *ccr4* (blue), *not1* (purple), and *pop2* (black) knock-downs as described in B-E. The levels of maternal transcripts were compared using relative RT-qPCR. The boxes (with upper and lower quartiles) indicate relative mRNA levels as a percentage of the control knock-down. N = 3 (combined 3 technical replicates from 3 independent experiments). Error bars = SD.

To measure the levels of *bcd* mRNA and other maternal transcripts (*osk*, *nos*) in each knock-down, and to analyze their correlation with the presence of P-bodies, we quantified the mRNA levels in knock-down background via relative RT-qPCR. The germline-specific transposon *HetA* was used to control for any reduction in germline volume resulting from each knock-down line. In the *not1* and *pop2* knock-downs, which are associated with loss of P-bodies, we observed a drastic reduction in *bcd* mRNA levels, along with significant, but smaller, decreases in *osk* and *nos* mRNA levels (Fig 2F). In contrast, the *pacman* and *ccr4* knock-downs exhibited little to no effect on the ovarian levels of *bcd* and other maternal transcripts (Fig 2F). Taken together, these results suggest that the stability of *bcd* and other maternal mRNAs is correlated with the presence of P-bodies; the knock-downs which resulted in the most severe loss of P-bodies also displayed the most striking reduction in maternal transcript levels.

### *bicoid* mRNA is a predicted target of miR-305

P-bodies have been previously implicated in the miRNA and siRNA pathways. Mammalian Argonaute proteins physically interact with GW182, a member of both the miRNA and P-body pathways (7). In a previous study, reporter mRNAs bearing miRNA target sites were localized to P-bodies in a miRNA-dependent manner (8). We therefore investigated whether miRNAs may mediate the translational repression of *bcd* mRNA in P-bodies. We began our analysis of the *bcd* 3’UTR by searching for predicted miRNA binding sites. Using the online miRNA site prediction tool TargetScan Fly (http://targetscan.org/fly-12) we identified a single, highly conserved miR-305 binding site in the first 100-nucleotides of the *bcd* 3’UTR, within a predicted single-stranded region known as Domain I (Fig 3A-B) (29). Interestingly, this binding site overlaps with *bcd*’s previously identified NRE motif (Fig 4A), alluding to the possibility of competition or cooperativity in the binding of Pum and miR-305. The NRE contains consensus motifs for Pum binding, and has been shown to regulate the stability of *bcd* mRNA during early embryogenesis (18).

**Fig 3.**
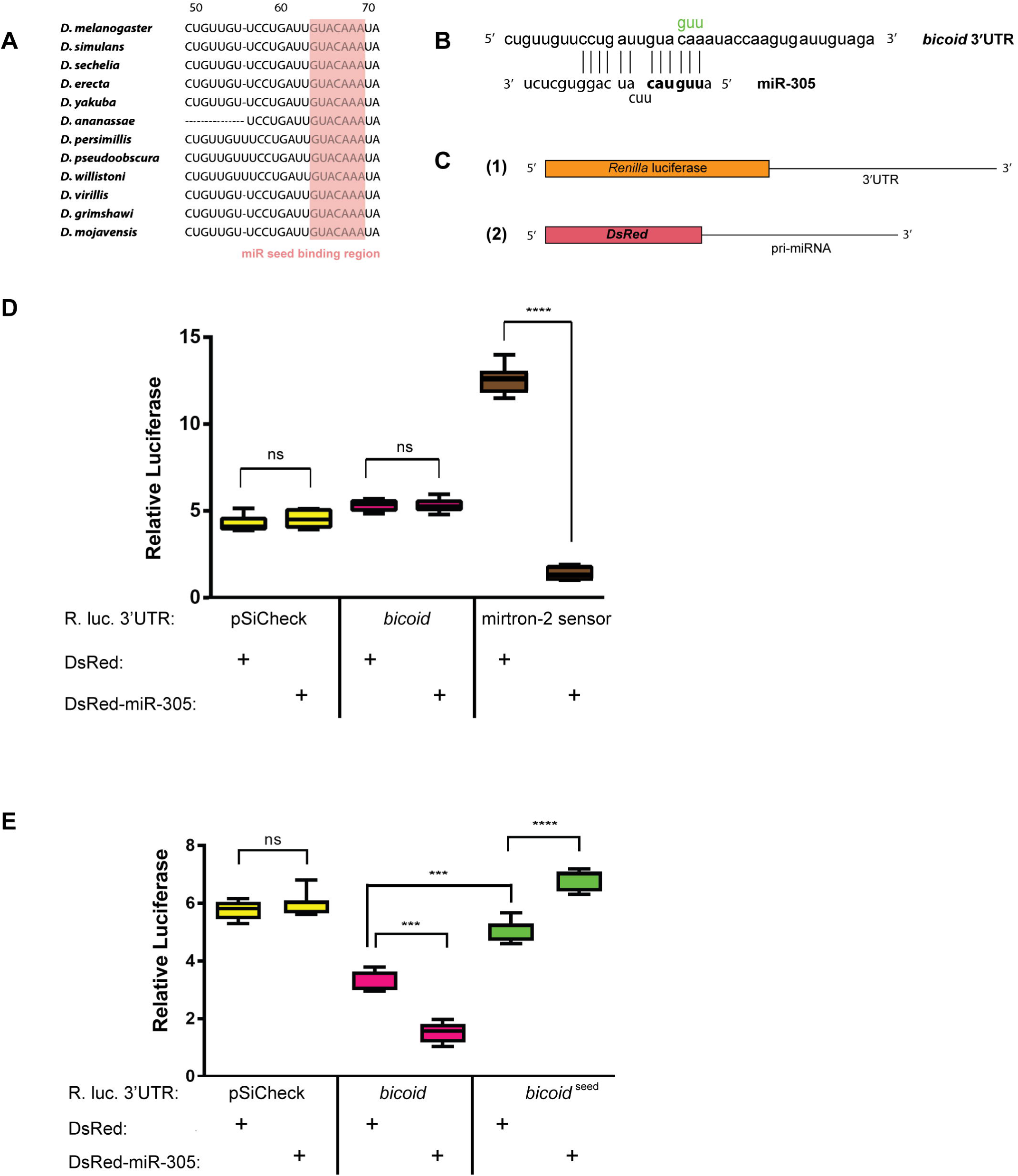
*bicoid* mRNA is a predicted target of miR-305. (A) Partial sequence alignment of the *Drosophilidae bcd* 3’UTR (TargetScan Fly). The predicted miR-305 seed-binding region is fully conserved among twelve *Drosophila* species (highlighted in red). (B) Predicted miR-305 base pairing with *bcd* 3’UTR; miR-305 seed region (bold) and “seed” mutations (green). (C) Reporter assay components: (1) pSiCheck 2 vector containing a *Renilla* luciferase reporter bearing the 3’UTR of interest (i.e. *bcd* 3’UTR) as well as a firefly luciferase gene to serve as an internal control for transfection efficiency, and (2) DsRed fluorescent reporter encoding a pri-miRNA of interest (e.g. miR-305) in its 3’UTR. These two vectors were co-transfected into *Drosophila* S2 cells, followed by lysis and luminescence measurements. (D) Relative luciferase levels (*Renilla*/firefly) indicate that miR-305 represses a *bcd* 3’UTR reporter (magenta) in *Drosophila* S2 cells, and this repression is abolished by a mutation in the miR-305 seed-binding region (*bicoid*^seed^ in green). Expression of the miRNA-containing vector was confirmed prior to cell lysis by visual inspection of live cells for RFP expression *** = p<0.001; ns = not significant. N = 3 (combined 4 technical replicates from each of 3 independent experiments). Error bars = SD.

**Fig 4.**
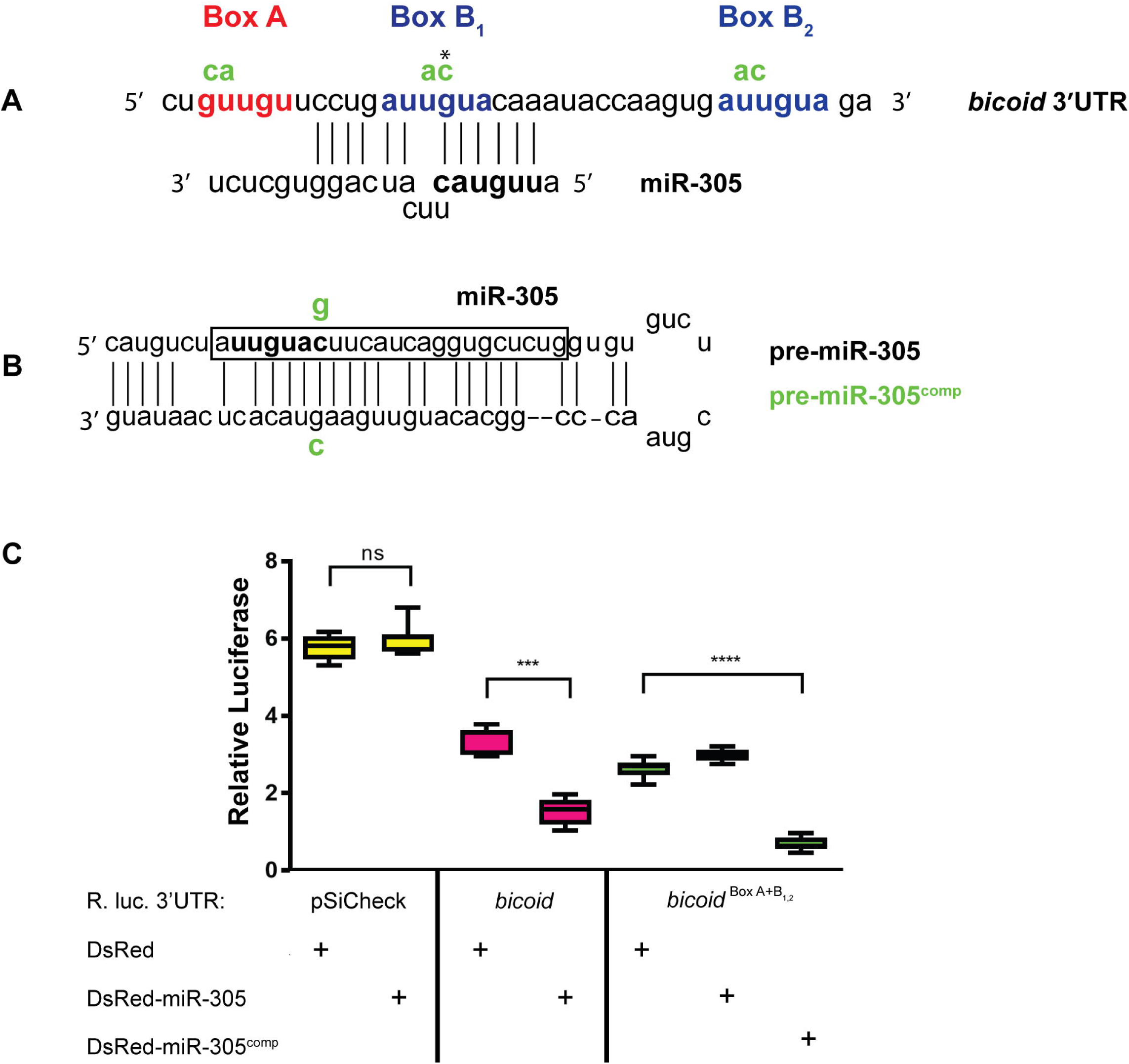
miR-305 represses a *bicoid* 3’UTR luciferase reporter independently of Pumilio binding sites. (A) Nanos Response Element (NRE) organization in the *bcd* 3’UTR: Box A (red) and Box B (blue) sequences; miR-305 seed region (bold), dinucleotide mutations (green). (B) The pre-miR-305 hairpin sequence with the mature miRNA (boxed). Compensatory mutant pre-miR-305 hairpin (bottom) with mutated nucleotides in seed region (green). (C) Relative luciferase (*Renilla*/firefly) levels indicate that the compensatory mutant miR-305 (miR-305^comp^) can repress the *bcd* 3’UTR reporter in the context of the triple mutant NRE sequence (*bicoid*^BoxA+B1,2^). **** = p<0.0001; *** = p<0.001; ns = not significant. N = 12 (combined 4 technical replicates from each of 3 independent experiments). Error bars = SD.

### miR-305 represses translation of a *bicoid* 3’UTR reporter mRNA in *D. melanogaster* S2 cells

To assess miR-305’s capability to repress translation via the *bcd* 3’UTR, we employed the DualGlo (Promega) luciferase assay in *D. melanogaster* late embryonic S2 cells (Fig 3C). To first establish the validity of the assay, we transiently transfected S2 cells with a luciferase-*bcd* 3’UTR reporter gene along with a fluorescent DsRed vector expressing miR-305 or mirtron-2. Co-transfection of the pSiCheck-*bcd* 3’UTR reporter with DsRed-mirtron-2 had no effect on luciferase levels, suggesting that the *bcd* 3’UTR reporter is not repressed due to off target effects of a miRNA not predicted to bind its sequence (Fig 3D). However, co-transfection of DsRed-miR-305 and pSiCheck-*bcd* 3’UTR resulted in ∼50% repression of the *Renilla* luciferase signal, as compared to the empty DsRed control, suggesting that the observed repression was a specific effect of miR-305 activity (Fig 3E). A trinucleotide mutation within the miR-305 seed-binding region in the *bcd* 3’UTR completely abolished miR-305 mediated repression, indicating that miR-305’s repressive effect is dependent on full complementarity with this region (Fig 3E). The presence of miR-305 did seem to increase the luciferase levels of the *bcd* seed mutant reporter (Fig. 3E, last two boxes); one possible explanation is an unknown target of miR-305 that is expressed in S2 cells and specifically modulates translation of the *Renilla* or firefly luciferase reporter genes.

### Pum-binding sites are not required for miR-305 mediated translational repression

The predicted miR-305 binding site is located within *bcd* mRNA’s previously identified NRE, a sequence motif bound by the translational regulators Nanos and Pum (Fig 4A). Several pieces of evidence from recent studies suggest a role for Pum as an accessory component of the miRNA machinery in different cellular contexts. First, sequence analysis of different mRNAs has demonstrated that Pum binding sites are significantly enriched in the vicinity of high-confidence miRNA binding sites (30). In human cells, Pum binding has been shown to enhance miRNA recognition of different target mRNAs (31, 32). In addition, the *C. elegans* Pum homolog, FBF, as well as mammalian PUM-2, physically associate with Ago1 and eEF1A to attenuate translation elongation (33).

The NRE motif was first identified in the *hunchback* 3’UTR and it is composed of two different sequence motifs termed Box A and Box B. The *hunchback* 3’UTR contains two Box A and two Box B sites (2 NREs), while the *bcd* 3’UTR contains “1.5” NREs, consisting of one Box A and two Box B sites (Fig 4A), each representing a potential binding site for one Pum molecule (34). *pum* mRNA is highly expressed in *Drosophila* S2 cells (http://flyatlas.org/atlas.cgi). In order to test Pum’s possible contribution to miR-305-mediated translational repression via the *bcd* 3’UTR, the Pum-binding sites were mutated separately, and in combination, followed by co-transfection with DsRed-miR-305 or an empty DsRed vector (Fig 4A). These introduced mutations were identical to those used previously in the *hunchback* NRE, which have been demonstrated to abolish binding by Pum (Fig 4A) (20).

Because Box B_1_ overlaps with the miR-305 seed complementary region, it is not possible to mutate this site and then attribute the translational de-repression to the loss of either miR-305 or Pum binding. To circumvent this issue, a compensatory mutation was introduced in the pre-miR-305 hairpin (termed miR-305^comp^), which restores its wild type partial complementarity to the Box B_1_ mutant (Fig 4B). Thus, this allows an assessment of miR-305’s ability to repress translation of the reporter in the absence of Pum binding to Box B_1_. A triple mutant NRE had no effect on miR-305’s ability to repress the pSiCheck-*bcd* 3’UTR reporter (Fig 4C), and neither did any single mutation (S1 Figure).

### miR-305 is expressed in egg chambers

The presence of miR-305 in unfertilized eggs was verified via RNA sequencing analysis, suggesting that the miRNA is maternally loaded (35). Our own absolute RT-qPCR assay for detection of mature miRNAs in ovary RNA lysates confirmed the presence of miR-305 and additional miRNAs (S2 Figure). miR-305 is encoded on the left arm of chromosome 2, clustered with miR-275 and the non-coding RNA CR43857 (Fig 5A). The *cuc^1^* allele abolishes the transcription of CR43857, as well as that of miR-275, 305 (23). We verified the loss of miR-305 expression in each mutant allele combination by RT-qPCR (S3 Figure). A small amount of pri-miR-305 signal was detected in the *cuc*^1^/miR-305 KO allele combination, consistent with previous results using these fly lines (23). To assess miR-305’s ability to translationally repress an mRNA target *in vivo*, we used a miRNA sensor transgene containing the GFP coding sequence, driven by the *ubiquitin* promoter and bearing two miR-305 binding sites in its 3’UTR (Fig 5B) (23). The miR-305 sensor was crossed into miR-305 knock-out (KO)/+, *Df(2L)*/+, and *cuc^1^*/+ heterozygous backgrounds. We observed modest de-repression in each genotype, indicated by the increased GFP fluorescence in the nurse cell nuclei compared to the miR-305 sensor/+ control, suggesting that miR-305 is active in the germline cells of the egg chamber (Fig 5C).

**Fig 5.**
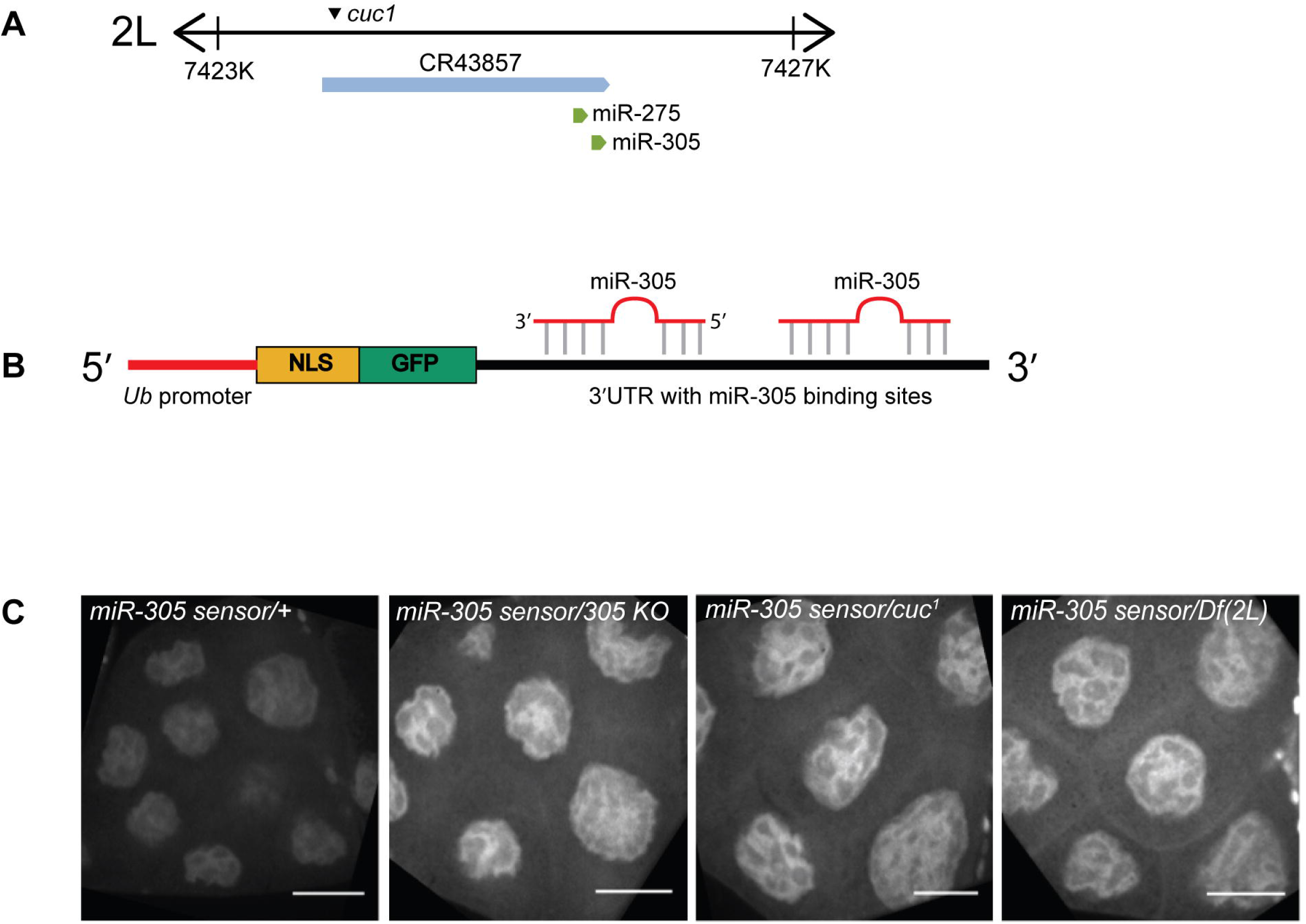
miR-305 is active in the egg chamber. (A) miR-275/305 are clustered within the CR43857 non-coding RNA located on chromosome 2L. *cuc^1^* is a null mutation in the promoter region of CR43857. Genomic positions are indicated; figure not drawn to scale. (B) Design of the miR-305 GFP sensor (23). The transgene encodes the GFP CDS, under control of the *ubiquitin* promoter, contains a nuclear localization sequence at the N-terminus, and optimal binding sites for miR-305 in the 3’UTR. (C) miR-305 GFP sensor expression in wild type, miR-305 KO, *cuc^1^*, and Df(2L) heterozygous mutant backgrounds. De-repression of the sensor in all mutant backgrounds was observed in the nurse cells as compared to the wild type background, as indicated by the higher intensity of GFP signal in the nuclei. Images were acquired with a 63X/NA = 1.4 oil objective, and each represent a single optical Z slice. Scale bar = 25 μm. Representative images were selected from 3 independent experiments.

### Transgenes for *in vivo bicoid* expression

To test the functionality of the miR-305 site *in vivo*, we cloned a *gfp-bcd* transgene for which expression is driven by the endogenous *bcd* promoter (Fig 6A). This transgene bears ∼200 nt of the *bcd* promoter sequence, previously demonstrated as sufficient for *bcd* expression *in vivo*. Embryos derived from *gfp-bcd* females exhibited anteriorly localized, nuclear GFP signal, demonstrating correct expression and localization of the transgene (Fig 6A). When this transgene was expressed in either miR-305 KO/+ or miR-305 homozygous KO genetic backgrounds, no GFP expression was observed, suggesting either that loss of miR-305 activity does not lead to premature translation of the *gfp-bcd* transgene, or that the amount of protein that is produced is below our detection limit (data not shown). We used the same *gfp-bcd* transgene to examine the functionality of the *bcd* NRE, by introducing it into a *pumilio* null background (*pum^1^* allele). There was no detectable GFP signal present in homozygous *pum^1^* egg chambers, suggesting that loss of Pum is not sufficient for premature *gfp-bcd* translation (data not shown).

**Fig 6.**
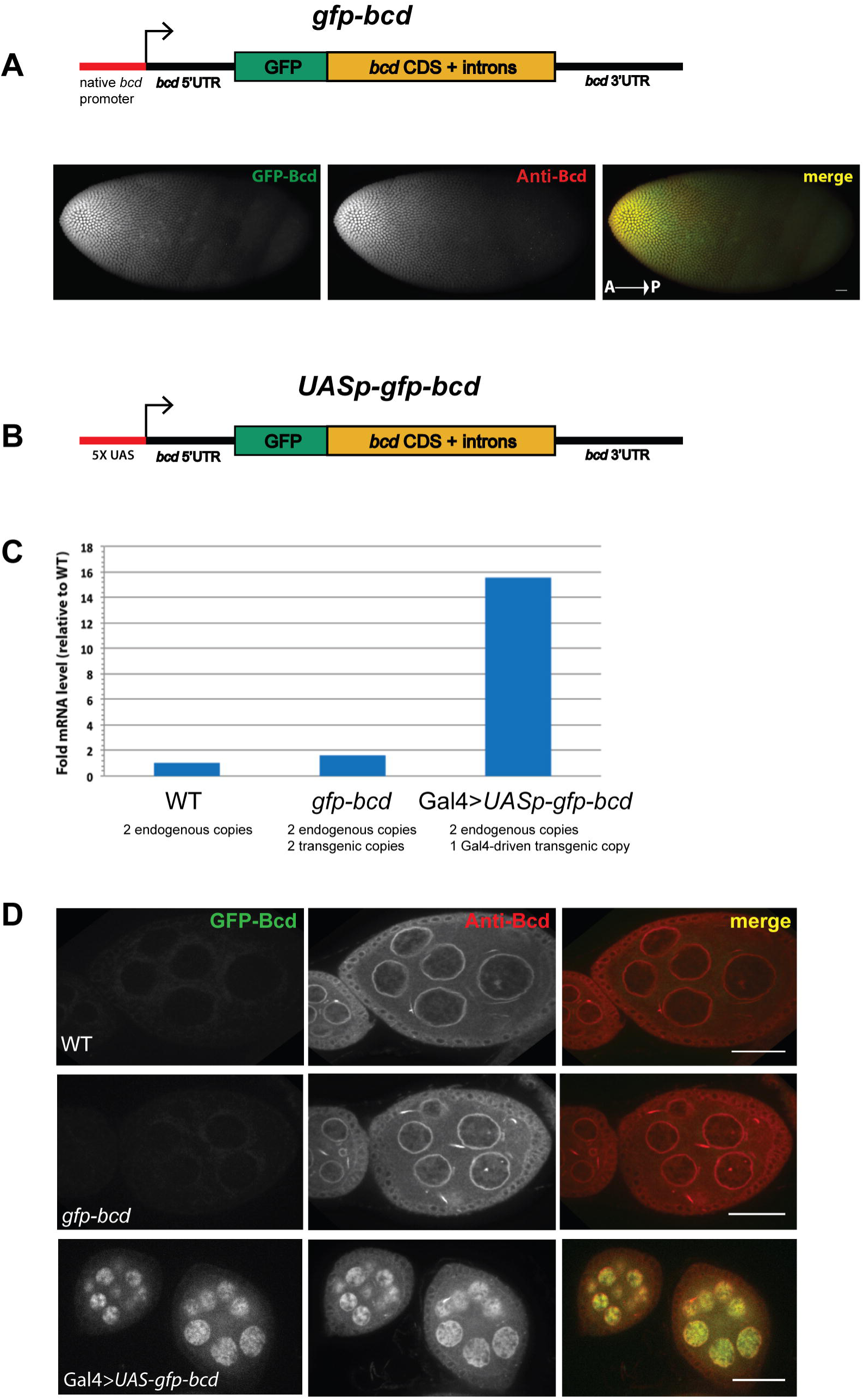
Expression of *gfp-bicoid* transgenes in the female germline. (A) *gfp-bcd* construct consists of the native *bcd* promoter and the *bcd* gene (CDS, introns, 5’ and 3’ UTRs), with an N-terminal GFP tag. The GFP-tagged Bcd protein correctly forms an A-P concentration gradient at the anterior of the embryo. Images were acquired with a 63X/NA = 1.4 oil objective, and are shown as stitched composite maximum intensity XY projections of 47 optical Z slices (Z step of 0.5 μm). Scale = 25 μm. The *gfp-bcd* transgene was crossed into a homozygous miR-305 KO background and ovaries examined for GFP expression using microscopy. There were no distinguishable changes in GFP levels between the miR-305 KO/KO and wild type ovaries. (B) *UASp-gfp-bcd* construct is identical to *gfp-bcd*, with the exception that the *bcd* promoter was replaced with a 5X UAS sequence. (C) *bcd* mRNA levels were quantified by relative RT-qPCR in ovaries isolated from the two transgene expressing lines. Bars represent samples combined from two independent experiments. (D) Ovaries isolated from the two transgenic lines were fixed and probed with a Bcd antibody (Cy5 label). Robust GFP and Bcd nuclear signals were observed only in the *UAS-gfp-bcd* early/mid stage egg chambers, accumulating predominantly in the nurse cell nuclei. Images were acquired with a 63X/NA = 1.4 oil objective, and are shown as maximum intensity XY projections of 3 optical Z slices (Z step of 0.5 μm). Scale bar = 25 μm.

### Transgenes for *in vivo bicoid* overexpression

Overexpressing *bcd* mRNA at high levels may oversaturate its endogenous repressive factors, allowing for the premature translation of *bcd* mRNA. To test this possibility, we cloned a new transgene identical to *gfp-bcd*, excepting the endogenous *bcd* promoter, which was replaced by 5X UASp sequence for Gal4-inducible expression (Fig 6B). Homozygous *gfp-bcd* ovaries expressed roughly twice the level of *bcd* mRNA compared to wild type ovaries, as expected, whereas the *UASp-gfp-bcd* line, driven by maternal Gal4, displayed ovarian *bcd* mRNA levels approximately 15-fold higher than those observed in wild type (Fig 6C). To determine if the high levels of *bcd* mRNA correlated with Bcd protein expression, ovaries isolated from each fly line were fixed and analyzed via immunofluorescence using a Bcd antibody. In *UASp-gfp-bcd* ovaries, robust expression of Bcd protein was detected as localized to nurse cell nuclei. Bcd was not detected in wild type or *gfp-bcd* expressing egg chambers, suggesting that just two additional copies of *bcd* gene are not sufficient to result in detectable premature levels of the protein (Fig 6D).

It is possible that in the absence of miR-305, the *gfp-bcd* transgene does produce a low, yet under our limit of detection, amount of Bcd protein. Therefore we used an alternative method to increase the baseline amount of GFP-Bcd protein and compare GFP expression levels between wild type and miR-305 seed mutant versions of the same *UASp-gfp-bcd* transgene (Fig 7A). Egg chambers of each genotype were examined at stages 3, 6, and 9 to quantify the pixel values of GFP fluorescence within nurse cell nuclei. Egg chambers from *UASp-gfp-bcd* seed mutant ovaries exhibited a statistically significant increase in GFP fluorescence levels at all stages, compared to those from the wild-type *UASp-gfp-bcd* transgene (Fig 7B). These results suggest that miR-305 binding may play a fine-tuning role in repressing *bcd* mRNA translation, which could not be examined using the *gfp-bcd* transgene expression in a miR-305 null background.

**Fig 7.**
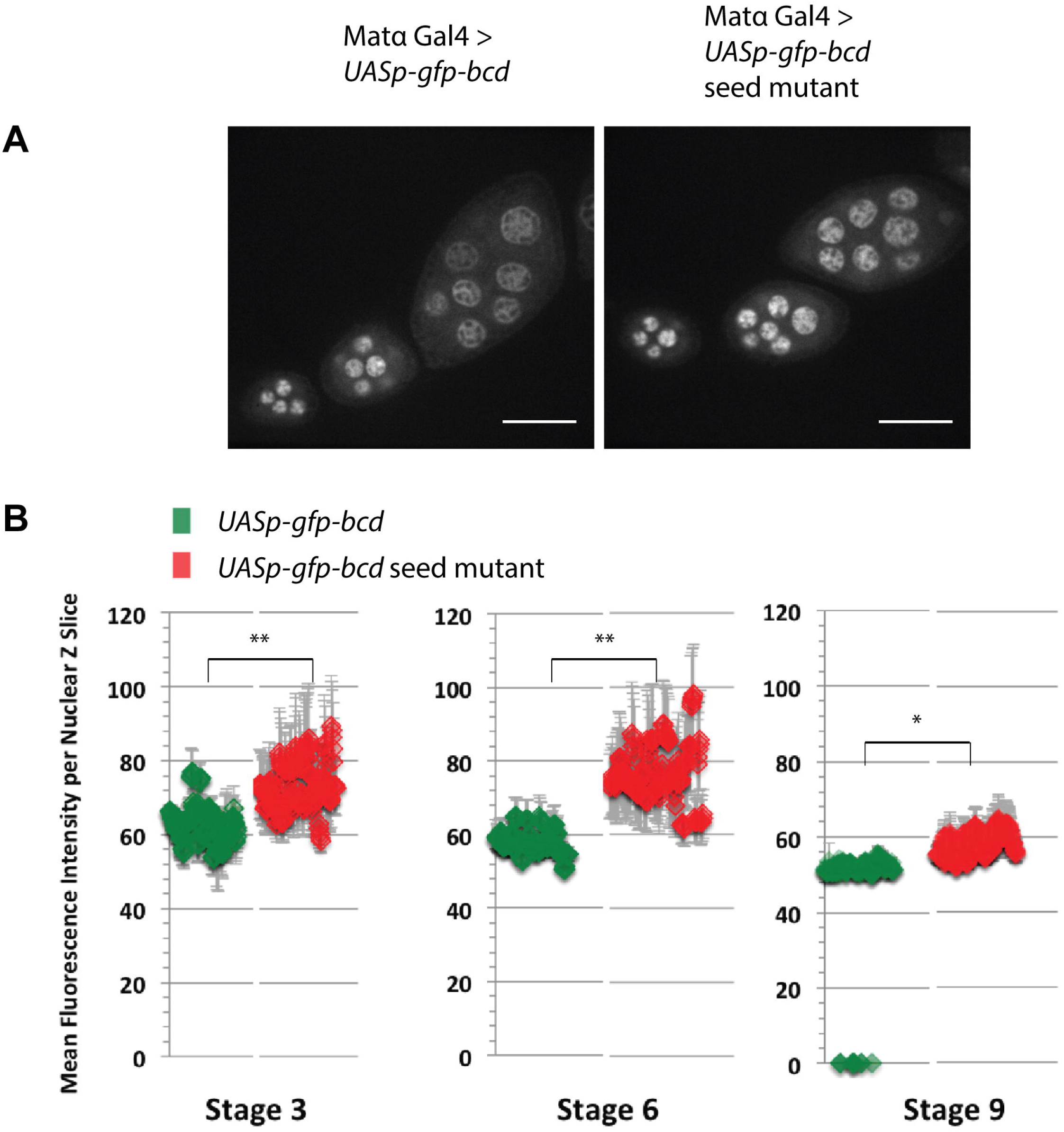
Quantitation of GFP expression from *UASp-gfp-bcd* transgenes. (A) GFP expression patterns in egg chambers isolated from wild type and seed mutant transgenes of *UASp-gfp-bcd*. GFP-Bcd is detected in the nuclei of nurse cells. Representative images were selected from the indicated genotypes. Images were acquired with a 63X/NA = 1.4 oil objective and are shown as maximum intensity XY projections of 3 optical Z slices (Z step of 0.5 μm). Scale bar = 25 μm. (B) Quantitation of nurse cell nuclear GFP fluorescence levels of each genotype was performed across 10 Z slices using ImageJ software, and plotted according to the indicated developmental stage. N = 3 (independent experiments). Error bars = SD. * = p<0.05; ** = p<0.005.

## DISCUSSION

Our understanding of gene regulation during oogenesis has been informed by decades of work on RNA-binding proteins, the transcripts’ cis-elements to which they bind, and how these interactions affect mRNA stability and localization. The ongoing study of post-transcriptional regulation by miRNAs is being integrated with this long accumulated understanding of gene regulation. Understanding translational regulation will require dissecting the mechanisms by which the miRNA pathway intersects with regulatory pathways involving different protein factors, as well as P-bodies.

### Links between mRNA P-body localization and translational control

We have provided evidence that the temporal control of *bcd* mRNA translation is regulated in part by the microRNA pathway, specifically by the activity of miR-305. We also found the stability of *bcd* mRNA to be positively regulated by P-bodies. This raises the question: to what degree do the separate mechanisms acting on the *bcd* 3’UTR interact or synergize, if at all? The *bcd* 3’UTR is highly structured according to enzymatic mapping, and organized into five separate domains that contain multiple stem-loops mediating RNA dimerization and localization (29, 36). Trans-acting proteins known to affect *bcd* mRNA localization include Staufen, Exuperantia, and Swallow, though their effects on its translation have not yet been examined. The contribution of these proteins to *bcd* mRNA localization is partially redundant, with loss of single factors, such as Staufen, displaying only modest effects on the localization process (37). To our knowledge, the function of the endogenous *bcd* NRE domain in its translational repression during oogenesis has not been previously explored.

Our data suggest that ovarian P-bodies act to stabilize *bcd* mRNA. This observation seems paradoxical, given the fact that P-bodies are sites of accumulation of the RNA decay machinery (28, 38). However, one recent study in support of our hypothesis demonstrated that in mutants of *D. melanogaster cup*, an eIF4E-binding protein and P-body component, *osk* mRNA levels are significantly decreased (39). This finding suggests that P-bodies can also play a role in stabilizing maternal transcripts. A possible explanation for this protective mechanism is that sequestration of *bcd* mRNA into highly organized P-bodies may “shield” the mRNA from exonucleases that would otherwise cause its degradation, or counteract the destabilizing effects of its short poly-A tail.

### Translational repression and activation

How is *bcd* mRNA’s translational repression relieved following egg activation? In *D. melanogaster*, egg activation is induced by mechanical pressure and rehydration of the mature egg chamber upon passage into the uterus (40, 41). A major cellular signal associated with this event is an increase in intracellular calcium levels. One possibility is that this initial signal causes a dynamic remodeling or breakdown of P-bodies in the oocyte, resulting in the free access of translational machinery to *bcd* mRNA. The calcipressin Sarah is required for egg activation; in *sarah* mutants, egg activation does not occur and *bcd* mRNA fails to be polyadenylated and translated during early stages of embryogenesis (42). Recent work from the Ilan Davis group has shown that egg activation correlates with *bcd* mRNA dissociation from Me31B foci, lending support to this hypothesis (6).

Another event concurrent with egg activation is the translation of Pan Gu Kinase (PNG). PNG is a major regulator of egg activation that mediates the translation of *cyclin B* (43). It is probable that PNG induces the translation of many additional maternal transcripts, both directly and indirectly, remodeling the proteome of the activated oocyte. One of these activated transcripts may be *bcd* itself, or a transcript whose protein product will activate *bcd* mRNA translation. It has also been previously demonstrated that a *D. melanogaster* poly-A polymerase, Wispy, is necessary for poly-A tail elongation of *bcd* mRNA following egg activation (44). Elucidating the order of events and their causal relationships will be critical to understanding the complete process of *bcd* mRNA translational activation.

### P-body composition and regulation of maternal mRNAs

Previous work suggests that P-bodies are dynamic structures, in contrast to more static cellular aggregates such as amyloid fibrils and stress granules (45). A possible mechanism for translational repression of *bcd* mRNA is via *bcd*’s sequestration in P-bodies, which exclude the translational machinery and ribosomes. Consistent with this possibility is our finding that overexpression of *bcd* in the female germline with the *UASp-gfp-bcd* transgene results in premature Bcd protein translation in the egg chamber. It would be interesting to analyze the sub-cellular distribution of the transgenic *bcd* mRNA, to determine whether it is present outside of P-bodies. It is possible that oversaturation of P-bodies with high levels of transcript results in access of *bcd* mRNA to the translational machinery.

This raises the question of how *bcd* mRNA is sequestered within P-bodies during oogenesis. Aggregation of many mRNA species and their trans-acting factors results in P-body formation. The observation that *bcd* mRNA is present within the P-body ‘core’ may simply be a consequence of *bcd* mRNA’s particular protein associations, rather than any active regulatory process. Overexpression of *bcd* mRNA may interfere with the stoichiometry of mRNA/protein interactions, shifting the equilibrium towards *bcd* mRNA being in higher abundance than usual outside of P-bodies. However, this outcome would also depend on whether P-body/*bcd* mRNA associated proteins are in large excess, and which protein factors are limiting in this interaction; adequately addressing this question requires the use of super-resolution microscopy techniques.

### Regulation of *bicoid* mRNA translational repression by miR-305

In *D. melanogaster* S2 cells, miR-305 exhibited robust repression of the *bcd* 3’UTR reporter, independently of its Pum-binding motifs. However, we were unable to detect any changes in the translational status of *bcd* mRNA in miR-305 KO egg chambers. We offer two possible explanations for this observation. First, miR-305 may not actually interact with the *bcd* 3’UTR in egg chambers due to a different accessibility to the mRNA in its native physiological setting. Second, miR-305 may only serve a modest, fine tuning role in repression of *bcd* mRNA translation, as opposed to functioning as a binary, all-or-nothing switch. There are many studies demonstrating that miRNAs mainly serve this type of minor, buffering role in gene expression control (46, 47). This more subtle role of miR-305 may also be partially concealed by redundant interactions among other trans-acting protein factors that contribute to the translational repression of *bcd* mRNA.

### RNAi screen for repressors of *bicoid* mRNA translation

An additional possibility is that other RNA-binding proteins, including known translational repressors, cooperatively or independently mediate the translational repression of *bcd* mRNA during oogenesis. We introduced the *gfp-bcd* transgene into RNAi backgrounds targeting different components of the miRNA pathway, P-bodies, and previously studied translational repressors (S4 Table; Genes screened by RNAi). Similarly to the *pum^1^* background, we did not observe an increase in GFP signal in any RNAi background relative to the control, suggesting that independent depletion of these genes does not alter the timing of *bcd* mRNA translation (data not shown). However, it is possible that removing combinations of these factors could result in premature Bcd expression, which would imply redundancy in the mechanism of its translational repression.

## Acknowledgments

We thank Eric Lai lab (Memorial Sloan Kettering Institute) for the reporter assay reagents, especially Dr. Diane Bortolamiol-Bécet for assistance with the *Drosophila* S2 cell experiments, and the Bratu lab members for stimulating discussion and comments on the manuscript.

---

**Fig S1. Single mutation analysis of *bicoid* Nanos Response Element via the *bicoid*-3’UTR reporter assay in *Drosophila* S2 cells.** The indicated NRE (B) *bicoid*^Box A^, (C) *bicoid*^Box B_1_^, and (D) *bicoid*^Box B_2_^ dinucleotide mutations were analyzed separately in luciferase reporter assays. Relative luciferase (*Renilla*/firefly) levels indicate that in each case, the single mutation has no effect on miR-305’s ability to repress the *bcd* 3’UTR reporter. **** = p<0.0001; *** = p<0.001; ns = not significant. N = 3 (combined 4 technical replicates from each of 3 independent experiments). Error bars = SD.

**Fig S2. Expression of mature miRNAs in ovary lysate.** (A) A stem-loop forming DNA primer, whose 3’ end is specific for a single mature miRNA, is added to a small RNA enriched lysate. To quantify levels of the miRNA, we employed a reverse transcription (RT) reaction followed by a conventional qPCR reaction. (B) The raw qPCR amplification curves for miR-305 (green) and other miRNAs. These curves were converted to histograms based on absolute quantitation methods, and presented as log(copy number) per 250 ng of small RNA enriched lysate.

**Fig S3. Expression levels of ovarian pri-miR-305 in mutant allele combinations.** The miR-275/305 KO, *cuc^1^*, and Df(2L) alleles were crossed into different trans-heterozygous combinations, and the ovarian pri-miR-305 levels measured by RT-qPCR (normalized to *rp49* mRNA). The expected loss of miR-305 expression was observed in all allele combinations. Each bar represents combined samples from two independent experiments.

