## Supplementary figures and images for "P-bodies and the miRNA pathway regulate translational repression of *bicoid* mRNA during *Drosophila melanogaster* oogenesis"

### Supplementary Materials

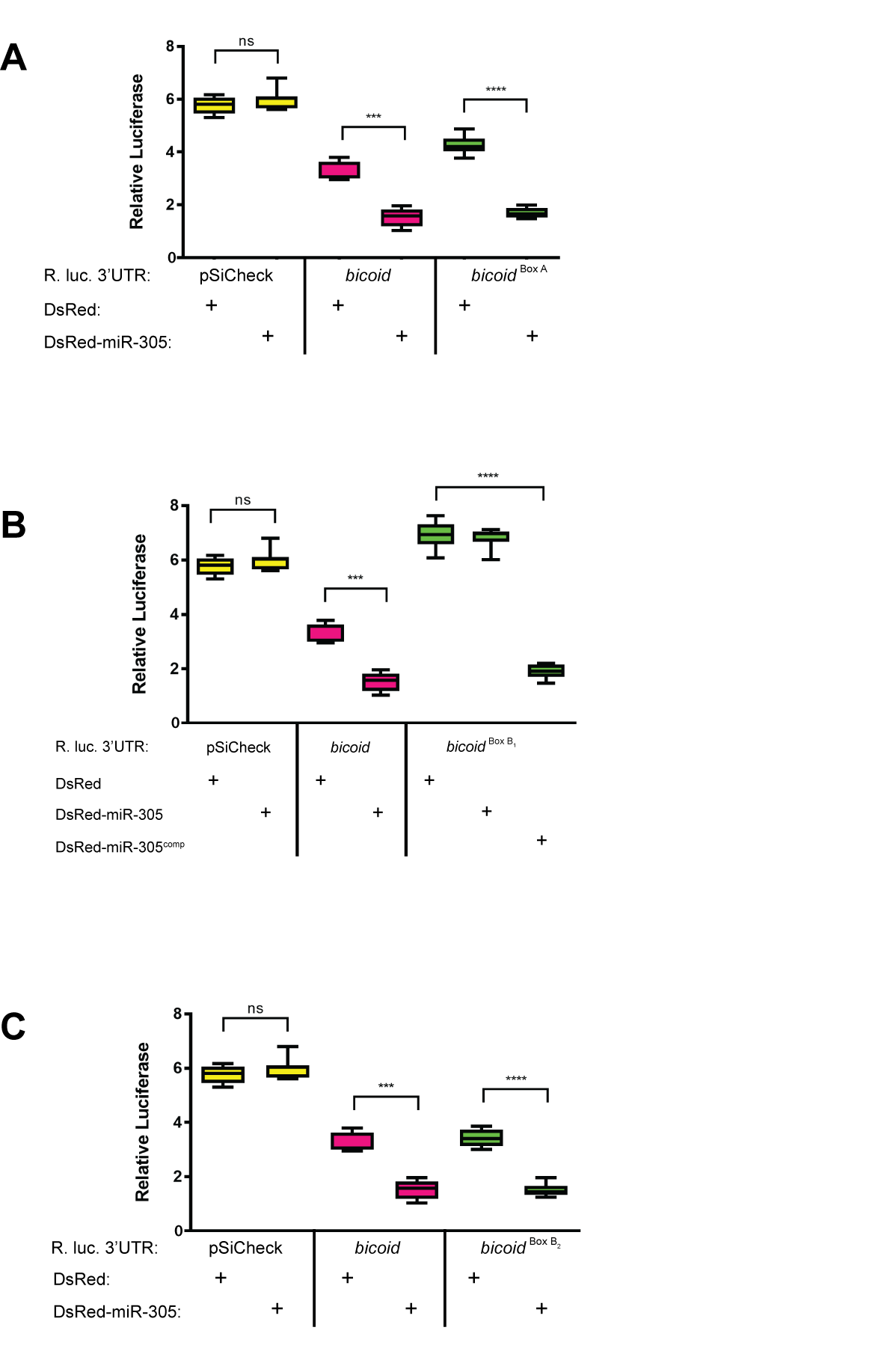

### Supplementary Materials

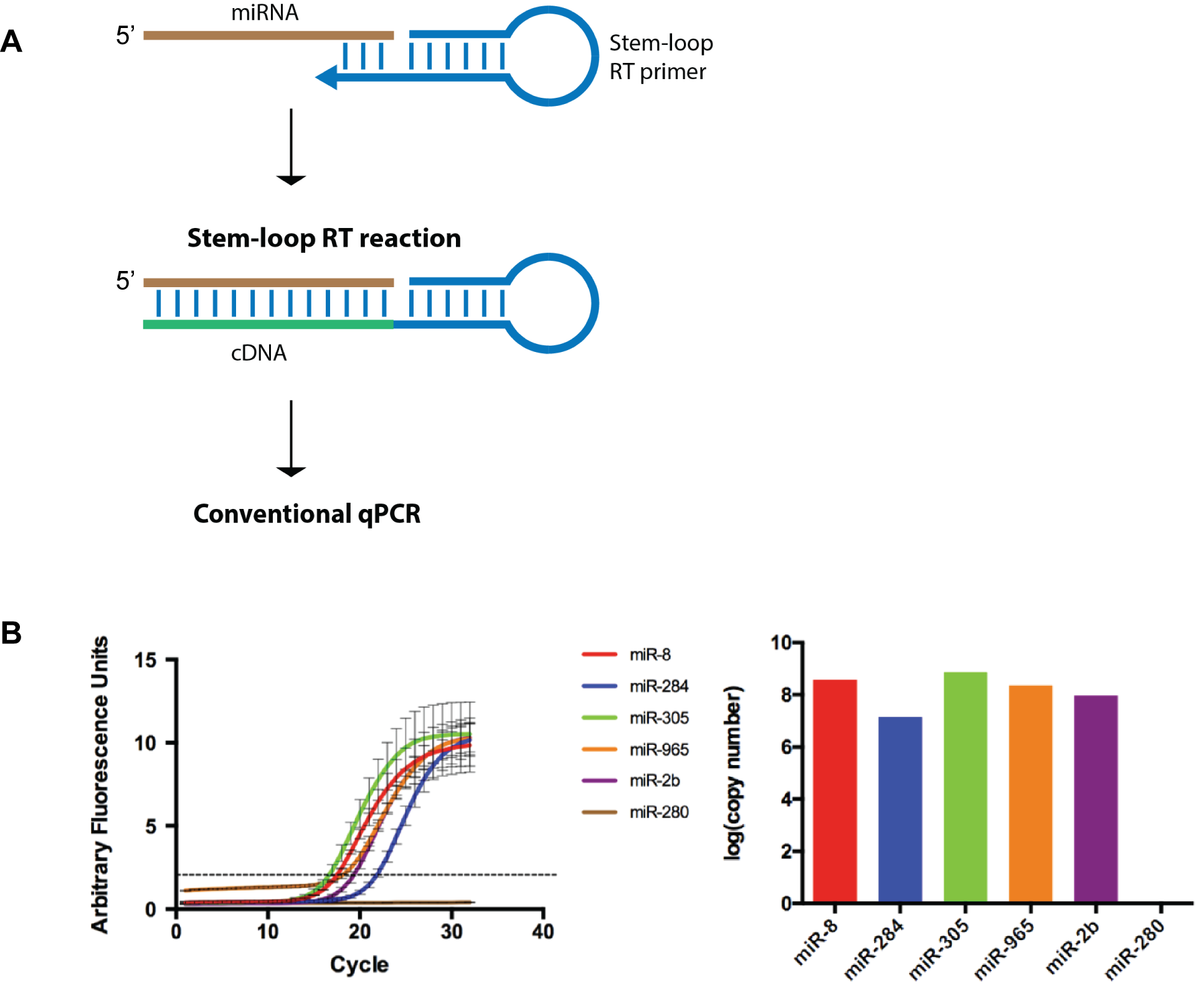

### Supplementary Materials

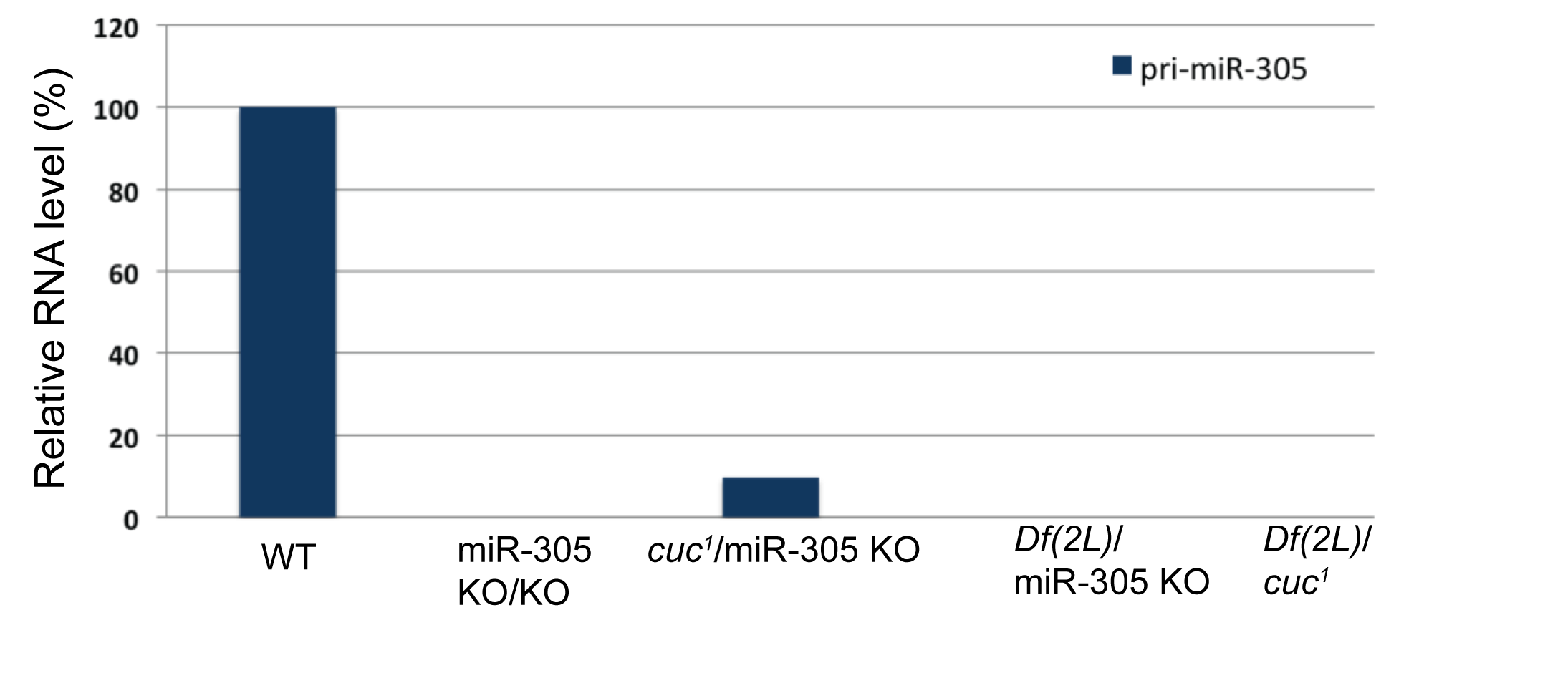
