## Supplementary Materials for "P-bodies and the miRNA pathway regulate translational repression of *bicoid* mRNA during *Drosophila melanogaster* oogenesis"

**TABLE S4**

**Oligo sequences for Gateway cloning**

| | |
|---|---|
| gfp-bcd fwd | caccatctcttcgctcatccctaaataacggcactctgcagatgcg |
| gfp-bcd rev | agttagtcacaatttacccgagtagagtagttcttatatattttcgt |
| gfp-bcd endogenous promoter fwd | caccgtctaa aatgtattgtagacgcttattg |

**Oligo sequences for qPCR**

| | |
|---|---|
| bcd PD44122 fwd | tgagcaccggaataagagcc |
| bcd PD44122 rev | ggcgttccgatggggattat |
| osk PP3222 fwd | atgaccatcatcgagagcaact |
| osk PP3222 rev | gtggctcagcaatatggcg |
| nos PP15723 fwd | ctcgtcggccactttgagtc |
| nos PP15723 rev | ctgtcggccagaaaagggaag |
| cdk9 PP5417 fwd | cgatgtccctgatggagaaac |
| cdk9 PP5417 rev | ccaattttcgccaccttttcgta |
| hetA fwd | cgcgcggaacccatcttcaga |
| hetA rev | cgccgcagtcgtttggtgagt |
| rp49 fwd | aagcccaagggtatcgacaacaga |
| rp49 rev | tcgacaatctccttgcgcttcttg |

**Oligo sequences for sequencing of gfp-bcd transgenes**

| | |
|---|---|
| gfp-bcd seq1 | atctcttcgctcatccctaaata |
| gfp-bcd seq2 | cgcatcgagctgaagggc |
| gfp-bcd seq3 | aattccgaaatcccttcgatttg |
| gfp-bcd seq4 | gcccaaccagatgcccaag |
| gfp-bcd seq5 | gcaggtccataatcaccagca |
| gfp-bcd seq6 | ccgatcgatgagatttgggag |
| gfp-bcd seq7 | cagtttgcctactgcttcaatta |

**Oligo sequences for restriction digest cloning**

| | |
|---|---|
| bicoid 3’UTR fwd | ttatcctcgagcctggatgagaggcgtgttagagagtttcattagc |
| bicoid 3’UTR rev | ttatcttgcggccgcgtagttagtcacaatttacccgagtagagtagttcttata |
| string (cdc25 ) 3’UTR fwd | ttatcctcgaggttgtgggatgatcgtgcagttcgttatctaag |
| string (cdc25 ) 3’UTR rev | ttatcttgcggccgccgtcgtgtattaatgtatatttaaaattgatgg |
| smaug 3’UTR fwd | ttatcctcgagaccccaatcacaacatcaactatttcattcacta |
| smaug 3’UTR rev | ttatcttgcggccgctatcccaactggccgtacaataggtttttatta |
| hunchback 3’UTR fwd | ttatcctcgaggttccccatcaccatcaccttgttattattattta |
| hunchback 3’UTR rev | ttatcttgcggccgcatattgaataattggatttatttgatttgatttcgttc |

**Oligo sequences for mutagenesis by PCR**

| | |
|---|---|
| bcd NRE Box A fwd | agagagtttcattagctttaggttaaccactcatgttcctgattgtaca |
| bcd NRE Box A rev | tgtacaatcaggaacatgagtggttaacctaaagctaatgaaactctct |
| bcd NRE Box B(1) fwd | ctttaggttaaccactgttgttcctgatactacaaataccaagtgattgtag |
| bcd NRE Box B(1) rev | ctacaatcacttggtatttgtagtatcaggaacaacagtggttaacctaaag |
| bcd NRE Box B(2) fwd | tgttgttcctgattgtacaaataccaagtgatactagatatctacgcgtag |
| bcd NRE Box B(2) rev | ctacgcgtagatatctagtatcacttggtatttgtacaatcaggaacaaca |
| bcd 305 seed fwd | cattagctttaggttaaccactgttgttcctgattgtagttttaccaagtgattgtagat |
| bcd 305 seed rev | atctacaatcacttggtaaaactacaatcaggaacaacagtggttaacctaaagctaatg |
| 305 comp c7g fwd | tgtctcccatgtctattgtagttcatcaggtgctc |
| 305 comp c7g rev | gagcacctgatgaactacaatagacatgggagaca |
| 305* comp g15c fwd | cgtaacccggcacatgttgaactacactcaatatga |
| 305* comp g15c rev | tcatattgagtgtagttcaacatgtgccgggttacg |
| smg 284 site 2 fwd | tacacataatttattttaaattaaaaggaacatatttcgtcaaccgtaactgcaccagag |
| smg 284 site 2 rev | ctctggtgcagttacggttgacgaaatatgttccttttaatttaaaataaattatgtgta |
| stg 965 site fwd | gttgtaaacttctgctaggtacaatttagcgttattatttgtttatttttatgtaatccg |
| stg 965 site rev | cggattacataaaaataaacaaataataacgctaaattgtacctagcagaagtttacaac |
| hb 8 site fwd | ctcagttctttctctgatattattctctgactttttttgttagttgaaagcgaattcgaat |
| hb 8 site rev | attcgaattcgctttcaactaacaaaaaaagtcagagaataatatcagagaaagaactgag |

**Oligo sequences for miRNA qPCR**

| | |
|---|---|
| miR-305 RT | gtcgtatccagtgcagggtccgaggtattcgcactggatacgaccagagc |
| miR-305 fwd | gcgattgtacttcatcaggtg |
| miR-284 RT | gtcgtatccagtgcagggtccgaggtattcgcactggatacgactgctg |
| miR-284 fwd | gtcagcaacttgattccagc |
| miR-8 RT | gtcgtatccagtgcagggtccgaggtattcgcactggatacgacgacat |
| miR-8 fwd | taatactgtcaggtaaagatgtc |
| miR-2b RT | gtcgtatccagtgcagggtccgaggtattcgcactggatacgacgctcct |
| miR-2b fwd | gctatcacagccagctttga |
| miR-965 RT | gtcgtatccagtgcagggtccgaggtattcgcactggatacgacaaggg |
| miR-965 fwd | taagcgtatagcttttcccc |
| miR-280 RT | gtcgtatccagtgcagggtccgaggtattcgcactggatacgactatcat |
| miR-280 fwd | gctgtatttacgttgcatatgaaatg |
